# A simple method to efficiently generate structural variation in plants

**DOI:** 10.1101/2024.12.20.629831

**Authors:** Lindsey L. Bechen, Naiyara Ahsan, Alefiyah Bahrainwala, Mary Gehring, PRV Satyaki

## Abstract

Phenotypic variation is essential for the selection of new traits of interest. Structural variants, consisting of deletions, duplications, inversions, and translocations, have greater potential for phenotypic consequences than single nucleotide variants. Pan-genome studies have highlighted the importance of structural variation in the evolution and selection of novel traits. Here, we describe a simple method to induce structural variation in plants. We demonstrate that a short period of growth on the topoisomerase II inhibitor etoposide induces heritable structural variation and altered phenotypes in *Arabidopsis thaliana* at high frequency. Using long-read sequencing and genetic analyses, we identified deletions and inversions underlying semi-dominant and recessive phenotypes. This method requires minimal resources, is potentially applicable to any plant species, and can replace irradiation as a source of induced large-effect structural variation.

## Introduction

Genomic structural variation (SV) – insertions, deletions, duplications, translocations and inversions – are an important source of genetic and phenotypic novelty. Structural variants create genetic novelty by multiple means, including fusing the coding regions of two genes, altering cis-regulatory environments and gene expression patterns, and by altering or suppressing recombination, among other mechanisms. In addition, deletions and duplications alter gene or chromatin dosage [1–3]. Novel structural variants have larger phenotypic effects per mutation than single nucleotide variants [4] and have been selected for when they underlie traits beneficial to the organism or desirable to agriculturists [5,6].

Recent pan-genome sequencing of multiple plant species has underscored the versatility and significance of structural variation to phenotypic variation, modern crop traits, and crop improvement. Lower seed sets in watermelons and grapes, important for the development of popular “seedless” varieties, have been linked to large inversions that cause meiotic defects [7,8]. SVs have also been linked to the domestication of broomcorn millet [9] and sorghum [10]. In pearl millets, gain of heat tolerance has been linked to SVs at multiple loci [11]. SVs are also associated with variation in immunity and plant defense responses [12–14]. In Arabidopsis, a multi-enzyme pathway synthesizes the glucosinolate family of defense compounds. SVs that either delete or fuse enzyme-encoding paralogs modify enzymatic pathways and are responsible for ecotype-specific variation in glucosinolates [15]. Studies of *Oryza sativa* provide numerous examples of how SVs influence phenotypes. Change in copy number of *VIL1* is associated with changes to flowering time and grain number variation [16]. In addition, duplications of the *KALA4* gene – a regulator of the anthocyanin biosynthesis pathway – created a novel cis-regulatory region that promoted ectopic expression of anthocyanins and caused a “black pericarp” phenotype [17]. In peaches, fruit flesh color around the stone and fruit shape are dependent on a deletion and a mega-base scale inversion [18]. Modern varieties of sweet corn were created by selecting for an inversion that creates a loss-of-function mutation in the *shrunken-2* gene, which encodes the first enzyme of the starch biosynthesis pathway [19,20]. In tomatoes, a duplication of a cytochrome P450 gene was linked to increased fruit weight [5].

Extant crop genetic diversity is unlikely to provide all possible SV diversity that could impact crop improvement. Induced structural variation can therefore provide novel germplasm for breeding programs, information that can be used for targeted CRISPR-mediated restructuring of crop genomes, and novel genetic variants for understanding fundamental aspects of plant biology. The most common method employed to induce random structural variation is exposure to ionizing radiation, including X-rays, gamma irradiation, and heavy ions [21–26]. Since the 1960’s, irradiation has been used to induce structural variation in model organisms and in numerous crops including, more recently, rapeseed [27], wheat [28], cotton [29], rice [23,30], poplar [22,31], soybean [32,33], and buckwheat [34]. Populations carrying such structural variation have been used for the isolation of mutations that abolish hybrid sterility in rice [35] and the creation of seeds differing in oil composition [21,27].

Although radiation-induced structural variation remains an important element of modern plant breeding, there are logistical challenges. In contrast to the induction of single nucleotide variants that researchers can conveniently pursue in their own labs using ethyl methanesulfonate (EMS) or sodium azide [36], the induction of structural variants requires access to radiation sources. Radiation sources are heavily regulated and access to nuclear reactors, particle accelerators, and other sources of radiation is a bottleneck in the creation of structural variant libraries [26]. Additionally, application of irradiation can be tedious. For example, the creation of a mutant population in poplar required irradiation of collected and dried pollen [22]. One alternative is the extended TAQing approach, in which a transgene expresses a restriction enzyme to induce conditional double-stranded DNA breaks, which then leads to shuffling of the plant genome [37]. However, this approach can be challenging as the commercial varieties of many crops are not easily transformed.

To surmount the current challenges of creating SVs in plants, we sought a simple method that could be applied to any plant species of interest to induce SVs with high frequency, drawing inspiration from the ubiquity and ease of EMS chemical mutagenesis to induce single nucleotide variants. One possible class of mutagens for SV induction is DNA topoisomerase II (Topo II) inhibitors. Topo II relaxes torsional stress from DNA supercoiling generated during DNA replication or transcription by transiently breaking both strands and then ligating them after passing a DNA segment through the break. Between strand breakage and ligation, Topo II is covalently linked to DNA via a tyrosine residue, forming a topoisomerase cleavage complex [38]. This complex is stabilized by the inhibitor etoposide. A collision between covalently-linked Topo II and DNA polymerases during DNA replication, or with RNA polymerases during transcription, leads to removal of the Topo II enzyme, which results in the generation of double-stranded breaks (DSBs) [39–42]. The imprecise repair of DSBs leads to genomic rearrangements and structural variation in mouse spermatocytes, fibroblasts, and in human cells [43–45]. Previously, it was shown that treatment with etoposide impacts genome stability and inhibits plant growth in *Arabidopsis thaliana*, *Allium cepa*, and *Lathyrus sativus* [46–49] and causes chromosomal fragmentation during meiosis in Arabidopsis [48]. However, its potential as a mutagen that can induce structural variation in plants has not been investigated. Here, we provide proof-of-principle that the chemotherapeutic drug etoposide efficiently generates novel genomic structural variation in *Arabidopsis thaliana*.

## Results

### Treatment with etoposide results in mutants with a variety of heritable phenotypes

To test whether etoposide can induce heritable structural variation in plants, we examined the effects of prolonged exposure to etoposide on Arabidopsis. Wild-type Col-0 seeds were germinated and grown on MS media supplemented with sucrose and differing concentrations of etoposide (0 µM (DMSO only), 20 µM, 40 µM, 80 µM, 160 µM, 320 µM, or 640 µM) for up to two weeks (Figure 1A). Plants grown on DMSO alone or 20 µM etoposide showed no observable growth defects (Figure 1A). Two-week-old seedlings grown on 40 µM etoposide exhibited gnarled leaves but no differences in size.

**Figure 1:**
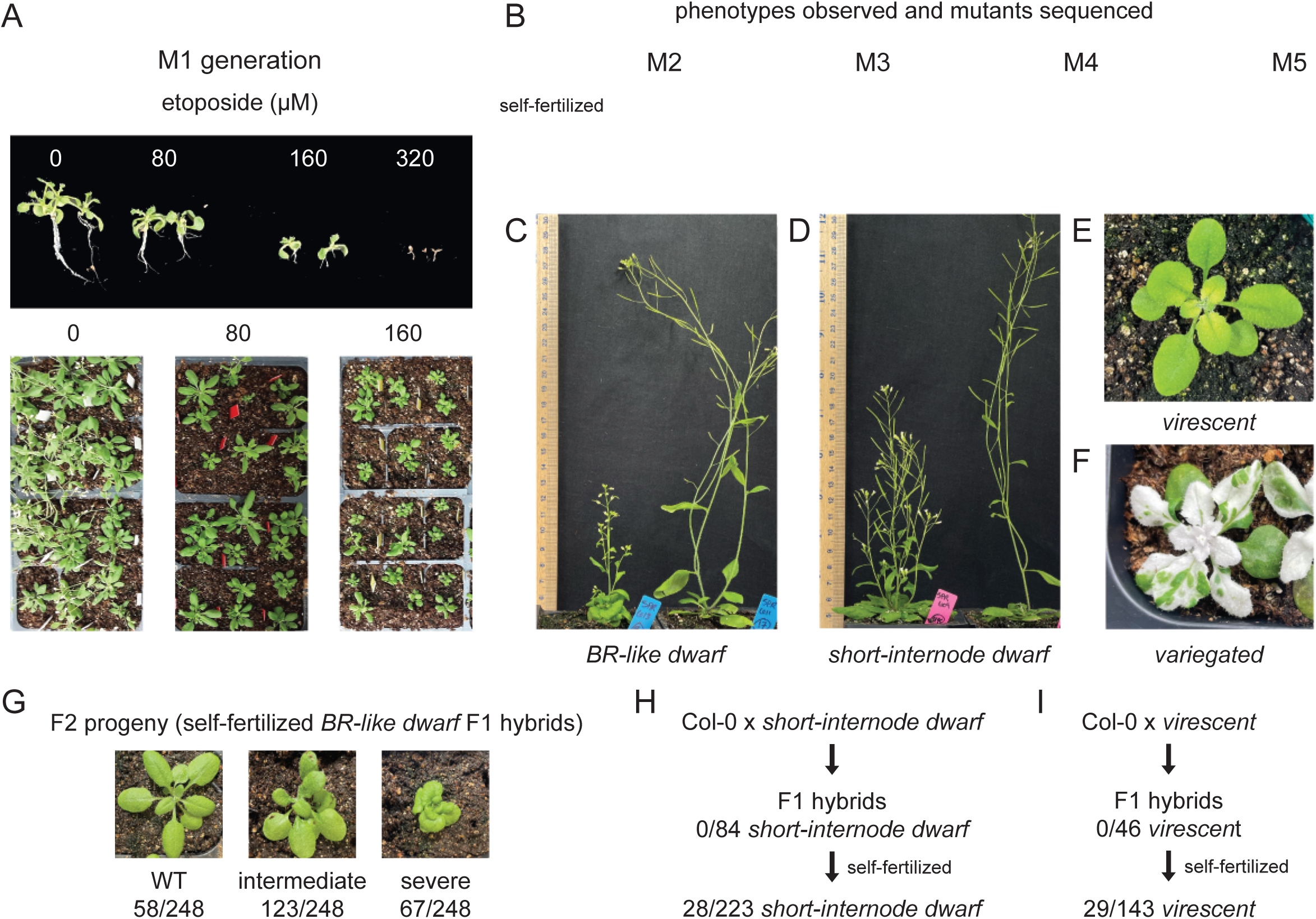
Etoposide induces novel heritable phenotypes. **(A)** Arabidopsis seeds germinated on increasing concentrations of etoposide show a dose-responsive growth defect (upper panel). Seedlings grown on 320 μM etoposide did not exhibit true leaves. This generation is referred to as M1. Growth defects persisted after transplantation to soil (lower panel). **(B)** M1 was self-fertilized to give rise to M2. Subsequent rounds of self-fertilization produced M3, M4, and M5 generations. A visual examination of M2 generation identified multiple phenotypes, including: **(C)** a *brassinosteroid-like dwarf* phenotype, **(D)** a fertile dwarf with short internodes, **(E)** a *virescent* phenotype with yellowish leaves, and **(F)** a *variegated* phenotype. **(G-I)** describe crosses to the wild-type and subsequent selfing to identify the nature of inheritance of the mutant phenotypes and the number of contributing loci. **(G)** The *BR-like dwarf* phenotype is incompletely dominant and displays three phenotypic classes. F2 segregation data suggests that this phenotype is caused by mutation at a single locus. **(H)** The *short-internode dwarf* phenotype is recessive and is likely caused by more than one locus. **(I)** The *virescent* phenotype is recessive and caused by a single locus.

**Figure S1:**
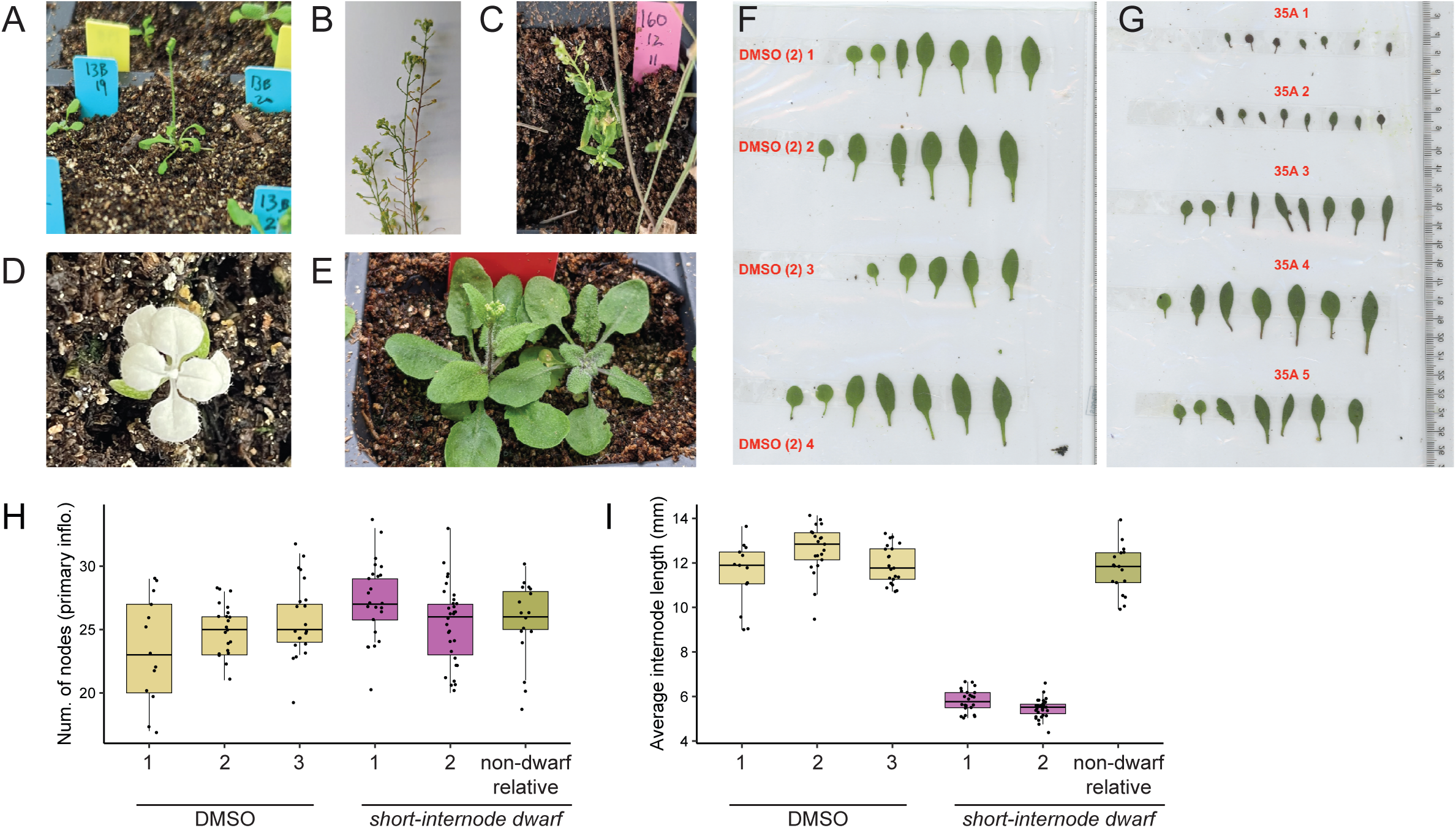
Additional phenotyping of progeny of etoposide-treated plants. Visual observation of M2 plants arising from different etoposide-treated M1 progenitors identified multiple lines with mutant phenotypes, including: early flowering **(A)**, sterility **(B)**, dwarfism **(C)**, variegation **(D)**, altered leaf shape **(E-G).** Lines with altered leaf shape or size were visually identified. These differences were confirmed by measuring leaf size (blade area, length, perimeter, width, and petiole length) and shape (cirularity) of leaves with the LeafJ plugin for ImageJ software. Leaves from multiple M2 siblings of line 35 are shown in **(G)**. 35A_1 and 35A_2 show a dramatic decrease in size parameters while 35A_3 shows an intermediate decrease. 35A_4 and 5 show no change compared to DMSO-treated lineages. **(H)** *short-internode dwarf* plants have the same number of nodes as DMSO-treated lineages and a genetic relative that lacks the phenotype, but **(I)** decreased distance between nodes, as measured on the primary inflorescence.

Seedlings grown on 80 µM etoposide showed marginally reduced growth (Figure 1A). Of the seedlings grown on 160 µM of etoposide, less than half (75/184) developed true leaves. In addition, these plants displayed stunted root development (Figure 1A).

Seedlings grown on 320 µM or 640 µM etoposide showed little root or shoot growth and high seedling lethality. Seedlings grown on DMSO only, 80 µM, and 160 µM etoposide (referred to as M1 plants) were transplanted to soil and compared at maturity. Those exposed to 160 µM of etoposide exhibited significantly more abnormal phenotypes than DMSO only or 80 µM etoposide plants, including loss of apical dominance, gnarled leaves, reduced plant size, seed abortion, and lower seed number at maturity (Figure 1A). To test if these abnormal plants produced progeny with mutant phenotypes, we scored for six visible phenotypes—dwarfism, loss of apical dominance, seed size or abortion, leaf shape or size, flowering time, and leaf pigmentation—among the progeny (M2 generation) of plants treated with 160 µM etoposide. Of M2 progeny derived from 42 different M1 parents, 29 lines exhibited at least one obvious visible phenotype (Figure 1B-F, Figure S1, Table S1). These phenotypes included: variegated or albino plants, altered flowering time, altered leaf shape and size, sterility or seed abortion, and dwarfism (Figure S1, Table S1). The large proportion of plants showing visible phenotypes suggested that etoposide could be an excellent mutagen for efficiently creating large-effect mutations.

To assess the heritability of the mutant phenotypes and to establish genotype-phenotype relationships, we closely examined four M1 lines with visible phenotypes: a dwarf line reminiscent of weak brassinosteroid-insensitive mutants (Figure 1C) [50], referred to as *brassinosteroid-like (BR-like) dwarf* (line 1A); a dwarf line with short internodes (line 34C), or *short-internode dwarf* (Figure 1D); a line with delayed greening (line 13B), termed *virescent* (Figure 1E); and a line with a *variegated* phenotype (line 5A) (Figure 1F). The semi-sterile *BR-like dwarf* line (Figure 1C) exhibited short, thick stems that bore fleshy leaves. In contrast, the *short-internode dwarf* mutant, though short in stature, was fertile, produced the same number of nodes on the primary inflorescence as WT plants (Figure S1H,I), and lacked the fleshy leaf phenotype observed in the *BR-like dwarf* (Figure 1D). In the *virescent* line, juvenile plants exhibited reduced chlorophyll pigmentation, whereas adult plants exhibited normal pigmentation (Figure 1E). *Variegated* mutants displayed light- and temperature-dependent variations in the proportions of green and white sectors on cotyledons and true leaves (Figure 1F).

**Figure S2:**
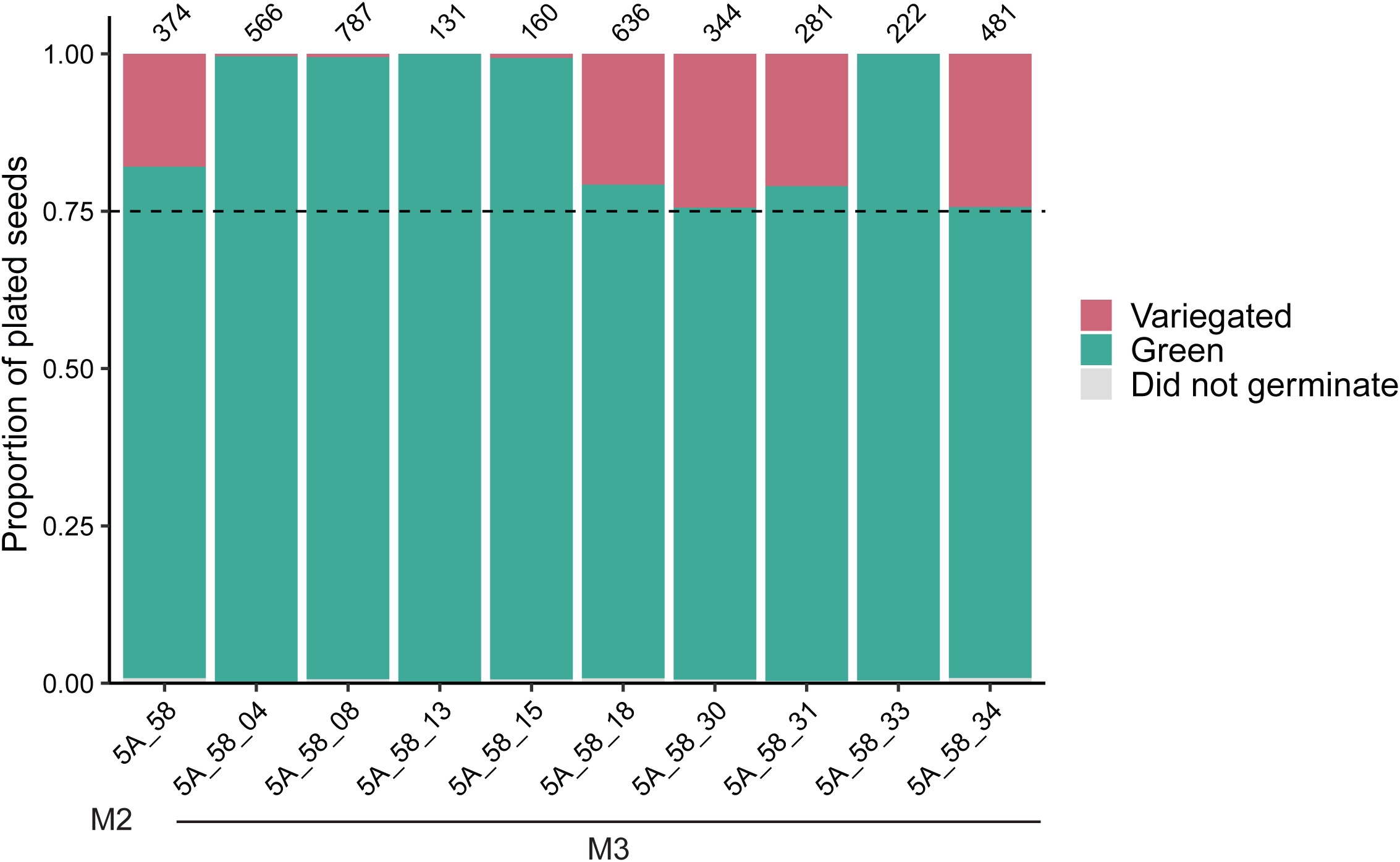
Segregation of the *variegated* phenotype from green parents. Green plants from variegated line 5A produce either approximately 25% variegated progeny or all green progeny. Number of seedlings assesed is annotated above each bar. M2 plant 5A_58 is the parent of the M3 progeny assayed.

The *BR-like dwarf, short-internode dwarf,* and *virescent* phenotypes were transmitted from self-fertilized parents to offspring for at least three additional monitored generations, indicating that the phenotype was true-breeding and the underlying mutant genotype was stable (Figure 1B). To assess whether these phenotypes were dominant or recessive, mutant lines were crossed to wild-type. The resultant F1 progeny from the *BR-like dwarf* exhibited intermediate dwarfing, suggesting that the phenotype was incompletely dominant (Figure 1G). The F1 progeny from the *short-internode dwarf* line and the *virescent* line displayed a wild-type phenotype, indicating that these mutant phenotypes were recessive (Figure 1H, I). *variegated* plants did not flower readily, so this phenotype was maintained as a heterozygous stock for the same number of generations. When self-fertilized, plants from this line either produced all green progeny or one-quarter variegated progeny, indicating that this phenotype is recessive (Figure S2).

To estimate the number of loci contributing to the *BR-like dwarf*, *short-internode dwarf*, and *virescent* phenotypes, we examined F2 progeny obtained by self-fertilizing the F1 plants. Of F2 progeny of *BR-like dwarf* plants with an intermediate dwarfing phenotype, 23.4% had a wild-type phenotype, 49.6% exhibited an intermediate phenotype, and 27.0% had a severe mutant phenotype (Figure 1G). This result suggested that a single incompletely dominant locus causes the *BR-like dwarf* phenotype (58:123:67 severe:intermediate:wild-type, H_0_=1:2:1 ratio, χ^2^=0.669; df=2, *p*>0.5). The *short-internode dwarf* phenotype was observed in 1/8 of progeny obtained by self-fertilizing F1 plants (28:195 *short-internode dwarf*:wild-type) (Figure 1H). This ratio is consistent with the recessive phenotype being caused by mutation in two linked loci, among other possibilities. For the *virescent* line, 20.9% of the F2 progeny displayed the mutant phenotype (Figure 1I). This is consistent with the phenotype being caused by a mutation at a single locus (29:114 *virescent*:wild-type, H_0_=1:3, χ^2^= 1.69; df=1, 0.25>*p*>0.1). These observations indicate that etoposide mutagenesis can efficiently create novel recessive and dominant phenotypes caused by large-effect alteration to one or more loci.

### RNA-Seq identifies genes that are associated with mutant phenotypes

To further characterize the four phenotypes studied in depth, we performed RNA sequencing of rosette leaves for M3 plants of *BR-like dwarf, short-internode dwarf, virescent,* and *variegated* lines (Table S2). As controls, we included genetic relatives that lacked the phenotype and M3 progeny of DMSO-treated plants. To obtain a broad overview of changes in gene expression, we performed gene set enrichment analysis (GSEA) for biological process gene ontology (GO) on the full ranked list of genes (ranked by log_2_ fold change) for each phenotypic line. We also identified differentially expressed genes for each phenotype, defined as genes with an adjusted *p*-value < 0.01 and a |log_2_(fold change)| > 1 (Tables S3-S5).

For the *BR-like dwarf* phenotype, a total of 309 genes were differentially expressed in *BR-like dwarf* plants compared to non-phenotypic plants (114 downregulated and 195 upregulated; Figure 2A, Table S3). The five most significant GO terms that were overrepresented among upregulated genes were: meristem development, anatomical structure formation involved in morphogenesis, response to auxin, plant organ formation, and post-embryonic plant morphogenesis (Figure S3A). *AS1* was a differentially expressed gene (log_2_FC = -2.62, adjusted *p*-value = 6.37 x 10^-16^), along with several genes it is known to interact with either directly or indirectly such as *KNAT1*, *KNAT2*, *KNAT6*, *STM*, *BOP1*, *BOP2*, and *LOB* (Figure 2A, Table S3) [51–54]. The phenotypes we observed in the *BR-like dwarf* plants were strikingly similar to those described for *as1* mutants, including a compact rosette with lobed and curled leaves [51,55].

**Figure 2:**
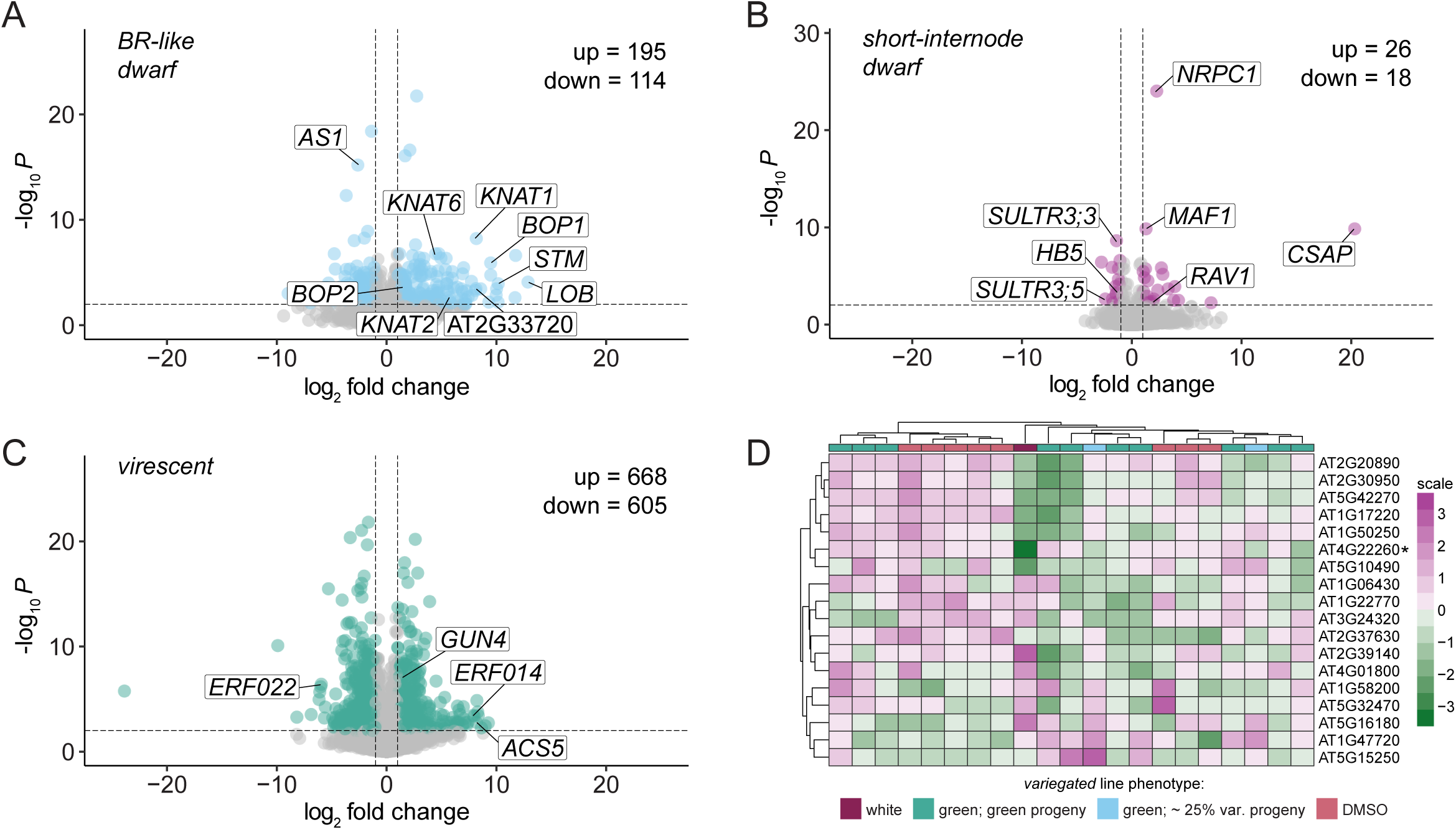
Etoposide-induced mutants exhibit changes in gene expression. Volcano plots summarizing differential expression analysis of phenotypic vs. non-phenotypic plants for **(A)** *BR-like dwarf*, **(B)** *short-internode dwarf*, and **(C)** *virescent* mutants. Significantly differentially expressed genes (|log_2_FC| > 1 and adjusted *p*-value < .01) are highlighted in (A) blue, (B) pink, and (C) green. Select genes are labeled in each volcano plot. **(D)** Heatmap of expression of genes known to be associated with variegation, in the *variegated* line. The most downregulated gene among them, *IMMUTANS,* is annotated with a *.

**Figure S3:**
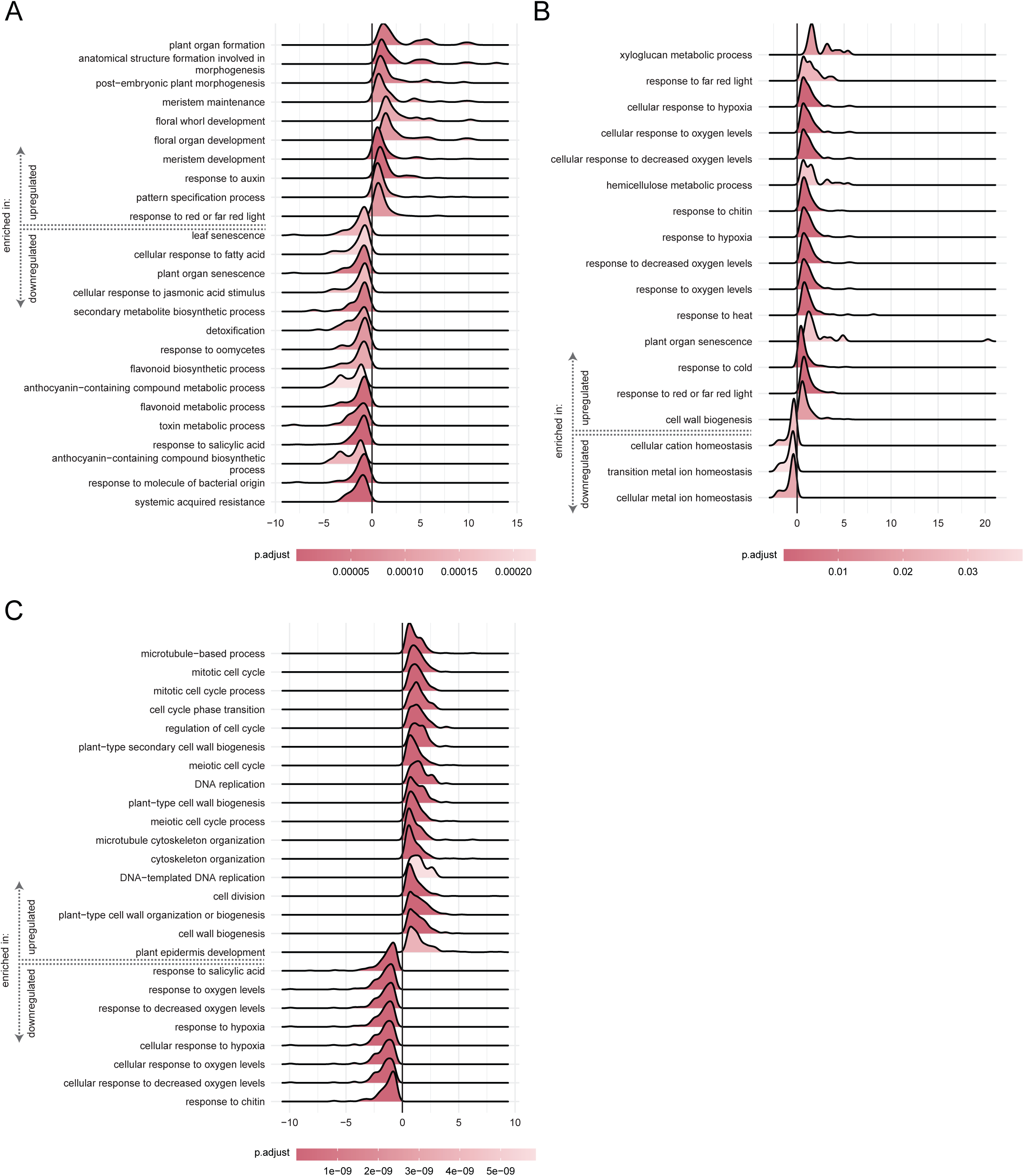
GO term enrichment for genes differentially expressed between plants exhibiting a phenotype and those without a phenotype. GSEA was used to identify enriched GO terms. Ridge plots depict significant terms for (A) *BR-like dwarf, (B) short-internode dwarf, and (C) virescent phenotypes*.

In contrast to *BR-like dwarf* plants, *short-internode dwarf* plants showed considerably fewer gene expression changes in leaves. A total of 44 genes (18 down and 26 up) were differentially expressed in *short-internode dwarf* leaves compared to control leaves (Figure 2B, Table S4). GSEA yielded primarily stress-associated GO terms (Figure S3B). Interestingly, both the largest subunit of RNA polymerase III, *NRPC1,* and the repressor of RNA polymerase III, *MAF1*, were upregulated. It is possible that *short-internode dwarf* plants exhibit changes in gene expression in other plant organs besides the leaf or perhaps exhibit a global change in gene expression that cannot be detected with standard RNA-seq approaches.

*Virescent* plants had more dysregulated genes than either dwarf line, with a total of 1273 differentially expressed genes (668 up and 605 down; Figure 2C, Table S5). The most significant GO terms overrepresented amongst genes with increased expression include cell division, plant-type cell wall organization/biogenesis, mitotic cell cycle, microtubule-based process, and meiotic cell cycle (Figure S3C). The most significantly overrepresented GO terms among genes with decreased expression include response to chitin, response to hypoxia, and response to oxygen levels (Figure S3C).

Interestingly, genes related to ethylene synthesis and signaling including *ERF022*, *ERF014*, and *ACS5* were among the most differentially expressed genes (Table S5). The mis-regulation of these genes might be associated with the *virescent* phenotype, as ethylene is involved in greening of etiolated seedlings after light exposure, and conversely degradation of chlorophyll during leaf senescence [56,57]. Other differentially expressed genes of note include several subunits of photosystem I and II and *GUN4* (Table S5).

Differential expression analysis could not reliably be conducted for the *variegated* line, as only one variegated plant was recovered. As an alternative strategy, we examined the expression of 18 genes known to be associated with variegation (Figure 2D). In the one variegated plant we recovered and performed RNA-seq on, the *IMMUTANS* (*IM*; *AT4G22260*) gene stood out as highly downregulated (Figure 2D). The sensitivity of the variegation to light and temperature observed in *variegated* plants is consistent with that displayed in *im* mutants [58,59].

### Etoposide treatment induces a spectrum of structural variation types

To characterize the molecular nature of etoposide-induced mutations, we employed Illumina short-read sequencing technology to sequence the genomes of 32 M2 or M3 progeny of 15 etoposide-treated M1 plants (Table S6). Sequenced plants included those with no scored mutant phenotype and those expressing a phenotype, including *BR-like dwarf*, *short-internode dwarf*, and *virescent,* among others. We also sequenced genetic relatives of these three mutants that displayed wild-type phenotypes (see Table S7 for relationships). To enhance the accuracy of our structural variation (SV) and other mutation calls, we also identified mutations segregating in our laboratory Col-0 population by sequencing four M2 plants descended from four control M1 lines treated only with DMSO.

As the inhibition of topoisomerase II activity increases double-stranded DNA breaks in the genome [60], the short-read data was used to test if the repair of these breaks results in increased single nucleotide variation (SNV). However, SNV analysis identified a comparable spectrum and number of SNVs in etoposide-treated and control lines (Figure 3A,B), suggesting that etoposide did not induce excess SNVs.

**Figure 3:**
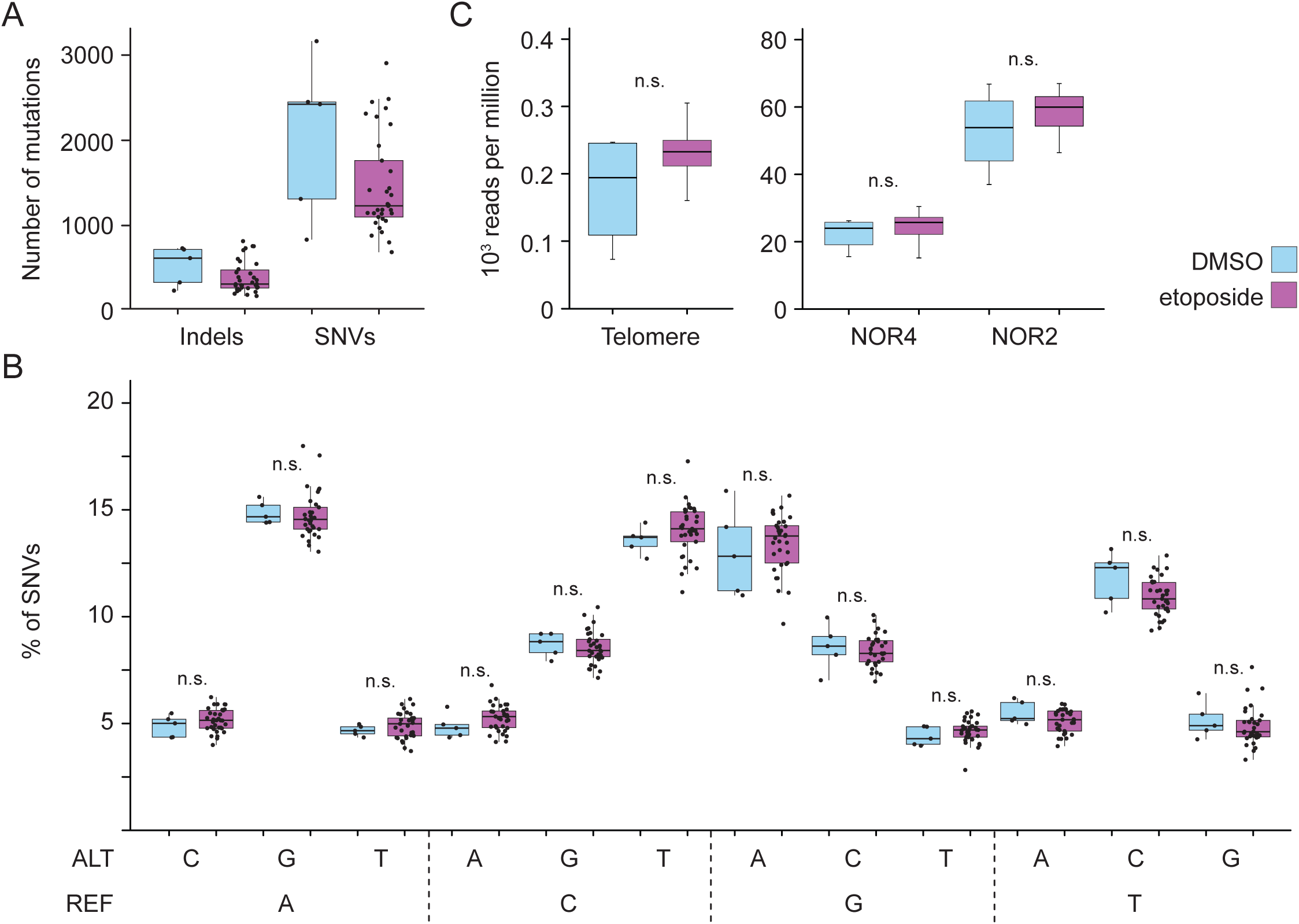
Impact of etoposide on SNVs and repetitive regions. **(A)** Total number of indels and single nucleotide variants (SNVs), genome-wide, identified in etoposide-treated (purple) and control lines (blue). **(B)** Box-plot describes the composition of nucleotide changes in progeny of plants exposed to etoposide and progeny of plants exposed to DMSO. All SNVs passing quality filters were included here. REF indicates reference allele and ALT represents the alternate SNV. Wilcoxon test was used to determine the significance of differences in SNV numbers between the progeny of etoposide and DMSO treated plant; n.s indicates *p >0.05*. **(C)** For Illumina short-read libraries of each control and etoposide-treated plant that we sequenced, total number of reads aligning to the telomere (CCCTAAA), NOR2, and NOR4 were normalized to total numbers of aligned reads in each library. Number of thousand reads, per million sequenced reads, for all control and etoposide treated libraries are represented in this boxplot. Wilcoxon test shows no statistical difference in the median value of read-depth over repetitive regions in control and etoposide-treated plants.

To test the potential for the induction of SVs in difficult-to-map repetitive regions, we measured read depth at rDNA and telomeres and assessed if etoposide treatment was linked to a higher degree of repeat copy number variation. While the number of control plants are insufficient to make a definitive statement, we found that the variation in read depth among etoposide-treated lines was comparable to that in control lines, suggesting that etoposide treatment likely did not trigger additional genomic instability in repetitive DNA (Figure 3C).

To identify etoposide-induced SVs, we first adapted a computational approach that uses genome-wide coverage to identify large segmental deletions or duplications [61,62]. Using this technique, we identified 10 duplications and 1 deletion that were between 100 kb and 1 Mb in length (Figure 4A, Table S8). The robustness of this analytical approach is underscored by detecting the same duplications in M2 siblings (Figure S4, Table S8). Next, we employed Lumpy Express [63] to identify structural variants (SVs) in mutant and control plants. This pipeline identified 2,224 variants that included inversions, deletions, duplications, and breakends, which were then filtered for high-confidence events. Breakends were excluded from further analyses because of the challenges of identifying their molecular nature. A high proportion of predicted SVs mapped to Nucleolar Organizer Regions (NOR), plastid genomes, and a small subset of other genomic loci; however these SVs were also found in control plants and read depth at NORs among etoposide-treated lines was comparable to that in control lines (Figure 3C), suggesting that these SVs represent mapping artifacts or naturally occurring SVs extant in wild-type plants. Based on these observations, we excluded SVs mapping to these regions from further analysis. To further filter for etoposide-independent SVs, those that were found in four or more etoposide-derived lineages were excluded, although we cannot exclude the possibility that these represent common fragile sites.

**Figure 4:**
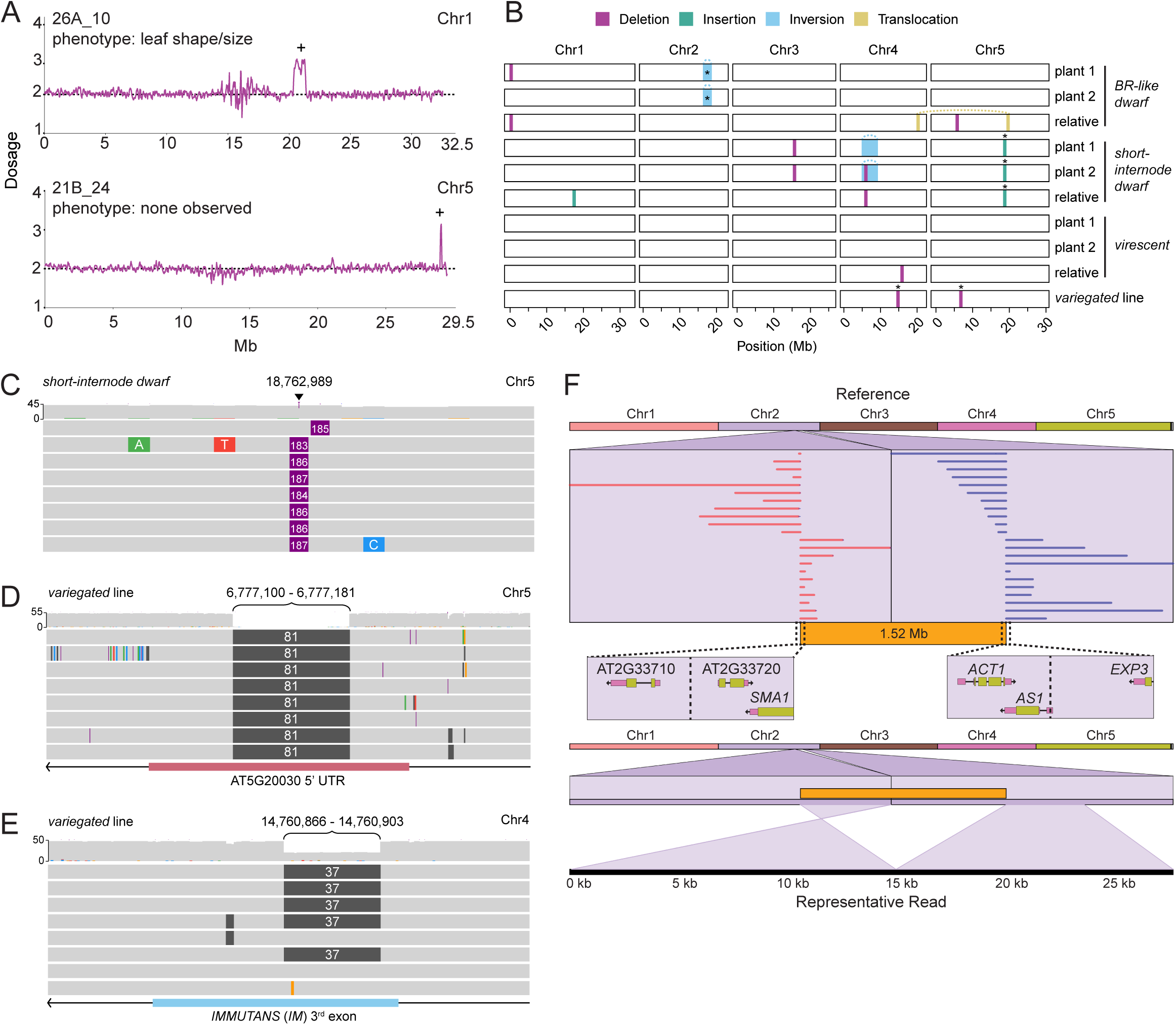
Whole genome sequencing identifies structural variation in progeny of etoposide treated plants. **(A)** Duplications in etoposide-treated lines as assessed by short-read sequencing. Dosage across the indicated chromosome is shown. Chromosomal segments with one extra copy are indicated with +. **(B)** Nanopore long-read sequencing identifies structural variation in lines that show the *BR-like dwarf*, *short-internode dwarf*, *virescent,* and *variegated* phenotypes. SVs verified via PCR are indicated with *. Putative causal mutations were identified by sequencing two plants with a phenotype and one relative without a phenotype. Only one mutant plant was sequenced for the *variegated* phenotype. This plant was green but produced ∼25% *variegated* progeny. Three PCR-verified mutations in (B) are described in (C-E). **(C)** A genome browser snapshot with reads identifying a 184 bp insertion on Chr5 in the *short-internode dwarf* line. **(D)** Reads in the genome browser snapshot indicate an 81 bp deletion on Chr5 in the *variegated* line. **(E)** Snapshot of long-read alignments to reference *IMMUTANS*, showing a 37 bp deletion in the third exon. Deleted sequence shown in grey and coverage is shown above read alignments. The deletion is heterozygous in the sequenced plant, which was green and produced ∼25% *variegated* progeny. **(F)** A ribbon plot with alignments indicating a 1.52 Mb chromosome 2 inversion in the *BR-like dwarf* line (top) and a detailed view of a representative read spanning the right breakend. The right breakend is in the 5’UTR of the *AS1* gene.

**Figure S4:**
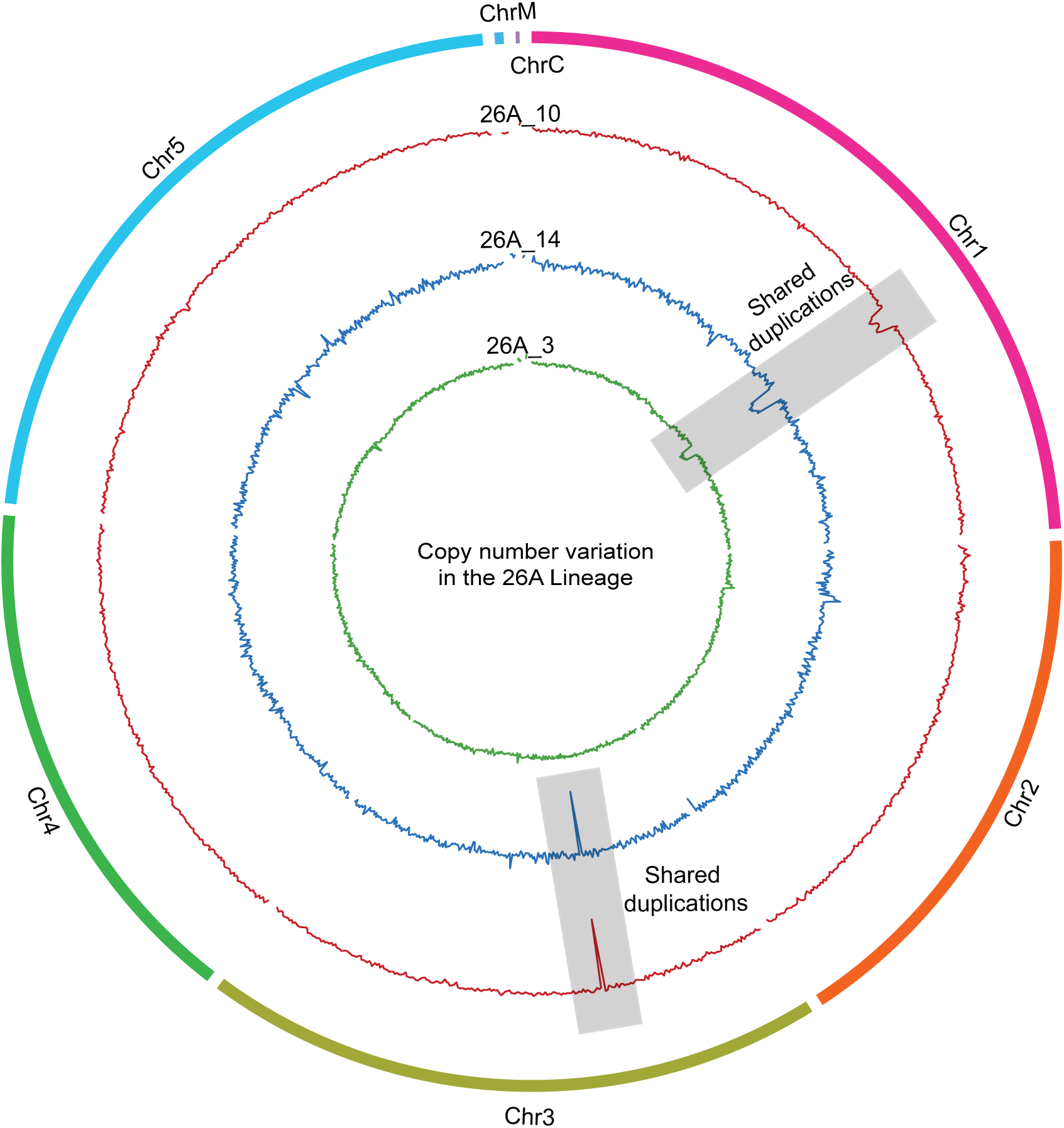
Duplications detected in the 26A lineage. Each track of the circos plot represents genome-wide read coverage analysis for an M2 sibling generated by selfing the 26A M1 mutant. Chromosome 1 duplications are shared by three M2 siblings while a Chromosome 3 duplication is shared by two of three siblings.

**Figure S5:**
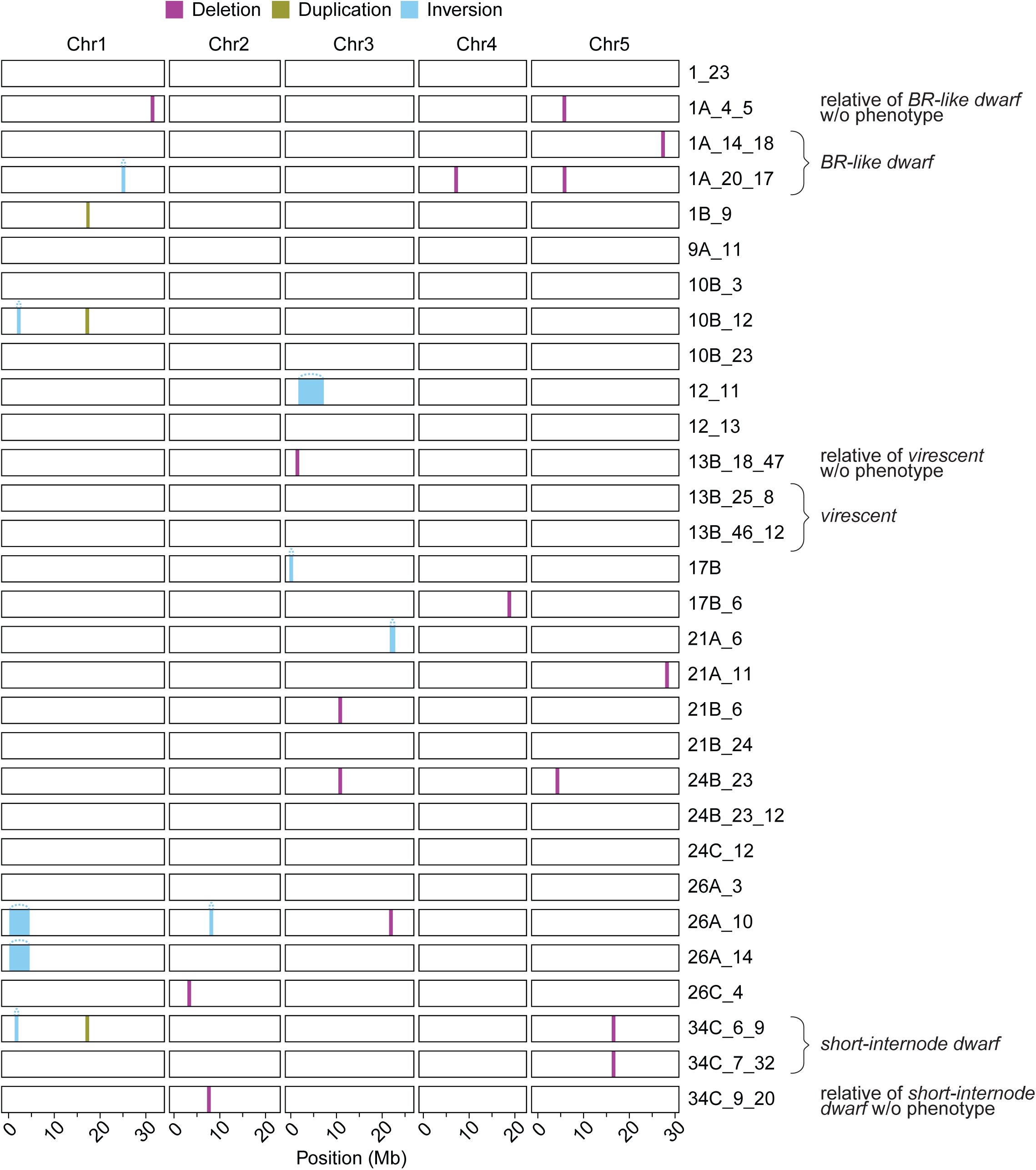
Structural variants identified by Lumpy Express in short-read data. A total of 27 SVs (16 deletions, 3 duplications, and 8 inversions) were identified. Eleven sequenced samples did not have any detected SVs.

After applying these stringent filters, we identified 27 unique SVs including 16 deletions and three duplications that ranged between 35 bp and 950 bp in size (Figure S5, Figure S6, Table S9). We also identified eight inversions that span between 124 bp and 4 Mb in length (Figure S5, Figure S7, Table S9). In sum, the coverage-based approach and Lumpy express analysis suggest that etoposide treatment induces structural variation in Arabidopsis.

Long-read sequencing can identify structural variants that are not easily resolved or cannot be identified by short-read sequencing [64]. Using Nanopore sequencing technology, we sequenced the genomes of two M3 plants with the *BR-like dwarf*, *short-internode dwarf,* or *virescent* phenotypes; an M3 relative of each corresponding line that lacked the phenotype; and a single M3 plant from the *variegated* line that was green but produced a proportion of variegated progeny (Table S10). To control for structural variation already present in our wild-type lab Col-0 stock compared to the reference Col-0 genome, we also sequenced two M2 plants that were the progeny of M1 plants grown on DMSO only and an untreated Col-0 plant. Reads were aligned to the Col-CEN v1.2 reference genome [65] and SVs were called using cuteSV [66,67]. Structural variants present in the DMSO and/or Col-0 controls, present in more than one M1 line, or with a quality less than 20 were discarded. In total we identified 12 SVs: seven deletions (ranging from 37 bp to 176 bp), two insertions (71 bp and 184 bp), two inversions (1.52 Mb and 3.54 Mb), and one translocation (Figure 4B; Table S11). We also chose to confirm several of the SVs via PCR (Figure S8): a 184 bp insertion with homology to an intergenic region of chromosome 2 present in the *short-internode dwarf* line (Figure 4C); a 37 bp and an 81 bp deletion present in the *variegated* line (Figure 4D,E); and a 1.52 Mb inversion present on chromosome 2 in the *BR-like dwarf* line (Figure 4F). Together, these data indicate that etoposide treatment generates a range of SV types and sizes.

**Figure S6:**
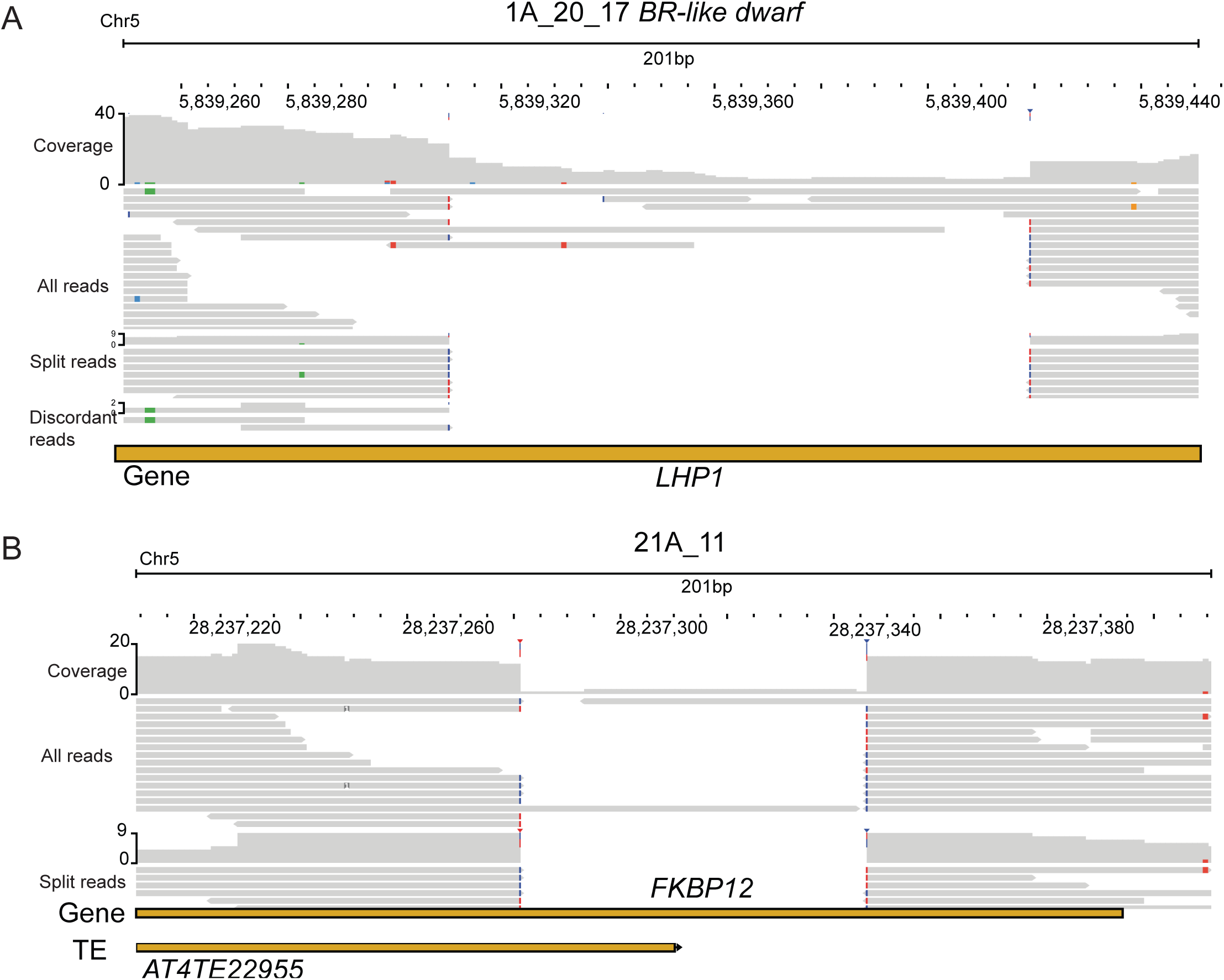
Examples of deletions in an etoposide-treated line identified by Lumpy Express using short reads. JBrowse snapshots of split reads and all reads overlapping *LHP1* **(A)** and *FKBP12* **(B)** identifies short deletions in two independent lines.

**Figure S7:**
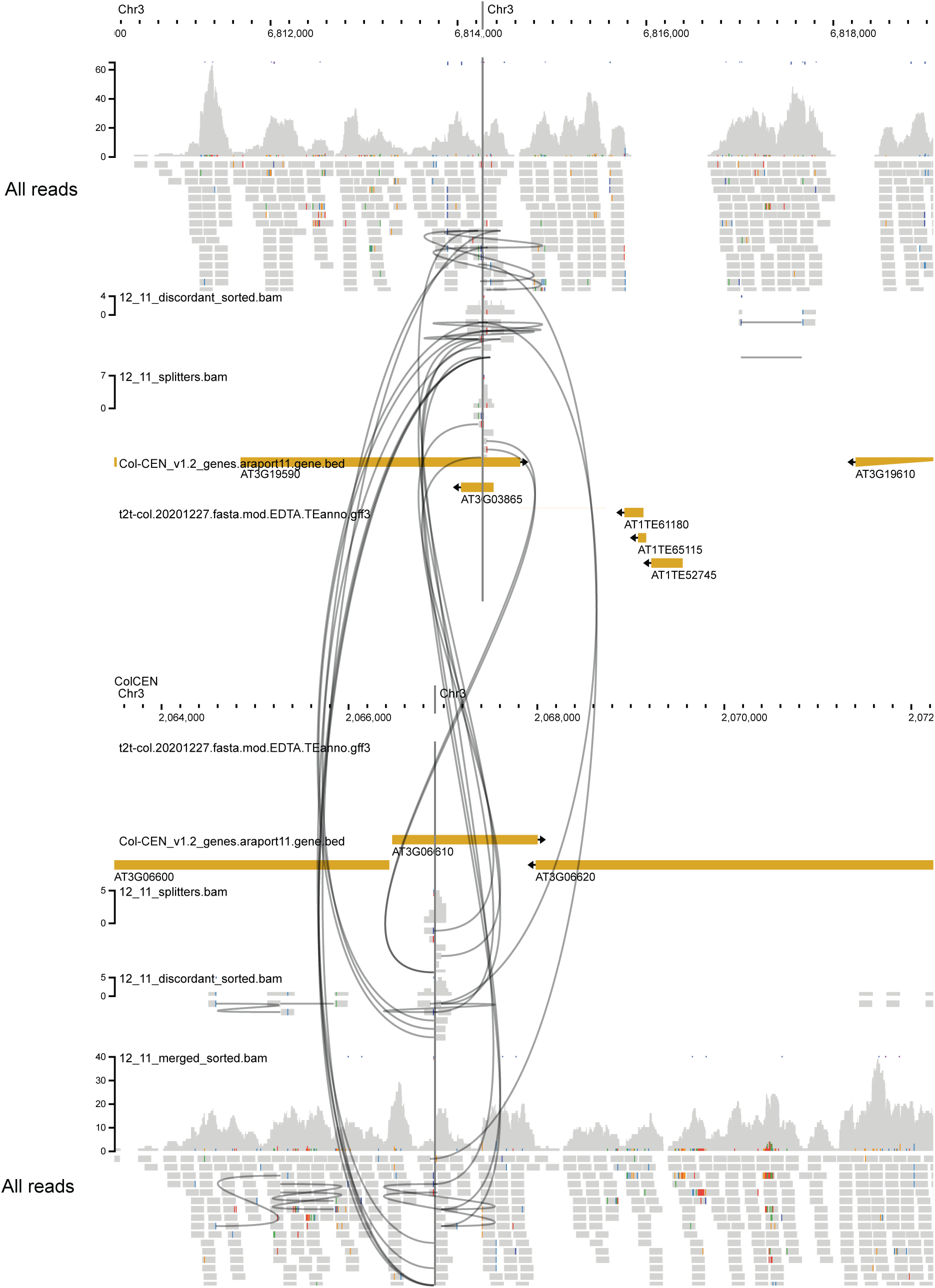
An example of an intrachromosomal inversion in an etoposide-treated line detected by Lumpy Express using short reads. Split reads and discordant reads indicate an intra-chromosomal inversion on Chr3. Black lines link split and discordant reads mapping to two different parts of the chromosome.

**Figure S8:**
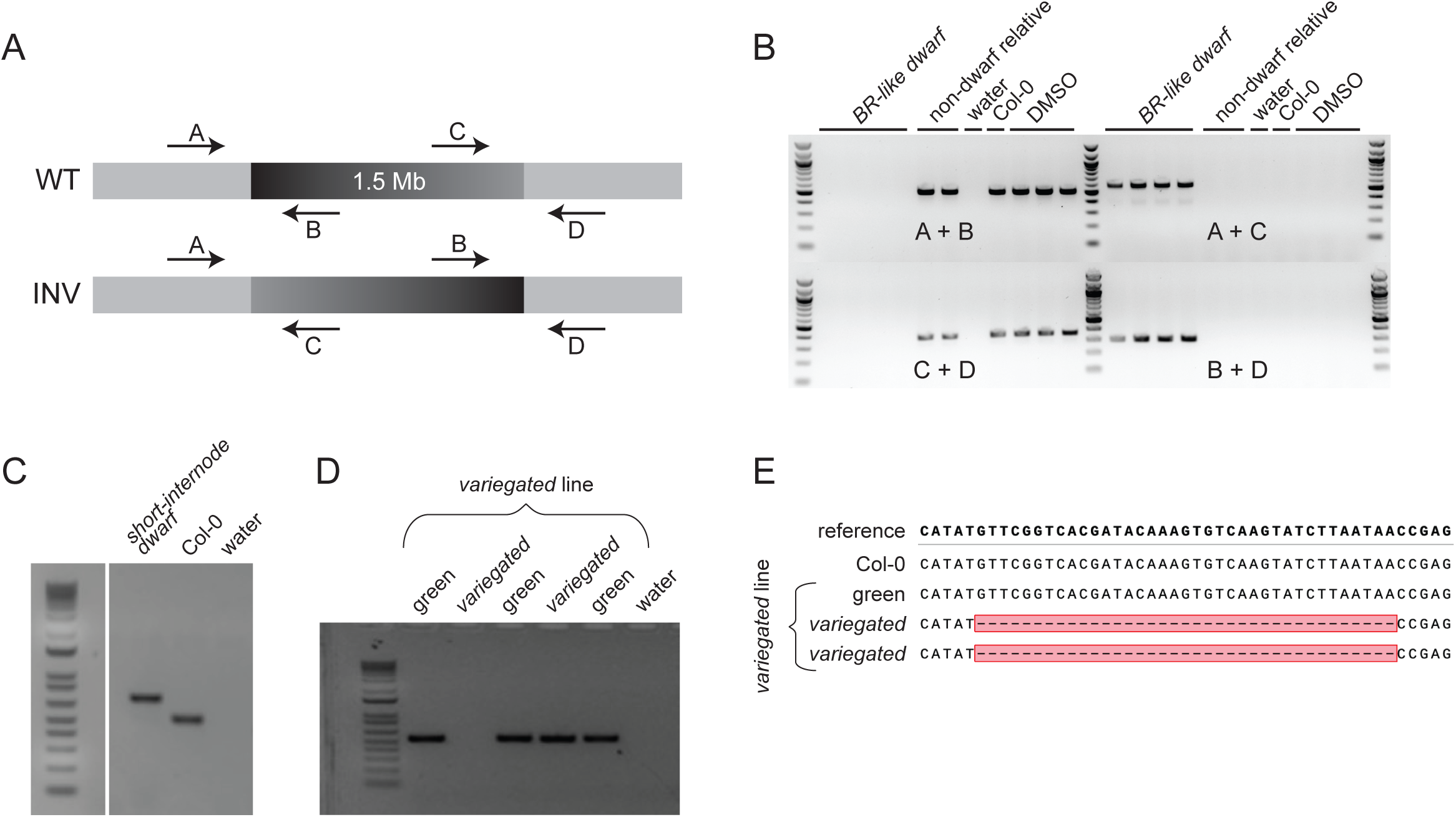
PCR verification of four strucutral variants identified via long-read sequencing. All plants sampled for PCR verification were relatives of the plants sequenced. **(A)** Diagram depicting primer design for verifcation of 1.52 Mb inversion identified in *BR-like dwarf* plants. **(B)** Image of gel electrophoresis of PCR products generated with different combinations of primers in (A); the inversion is present only in plants exhibiting the *BR-like dwarf* phenotype. **(C)** Gel image for verification of 184 bp insertion identified in *short-internode dwarf* plants. Primers were designed to flank the insertion. **(D)** Gel image for verification of 81 bp deletion identified in the plant sequenced from the *variegated* line. Primers were designed to flank the deletion; all plants tested are homozygous for the deletion, except for one plant (lane 3) for which the PCR failed. **(E)** Alignment of sequenced PCR products to *IMMUTANS* reference sequence. Primers were designed to flank the deletion. Individuals in the line exhibiting the *variegated* phenotype are homozygous for the deletion while green plants are either heterozygous or homozygous for the WT allele.

**Figure S9:**
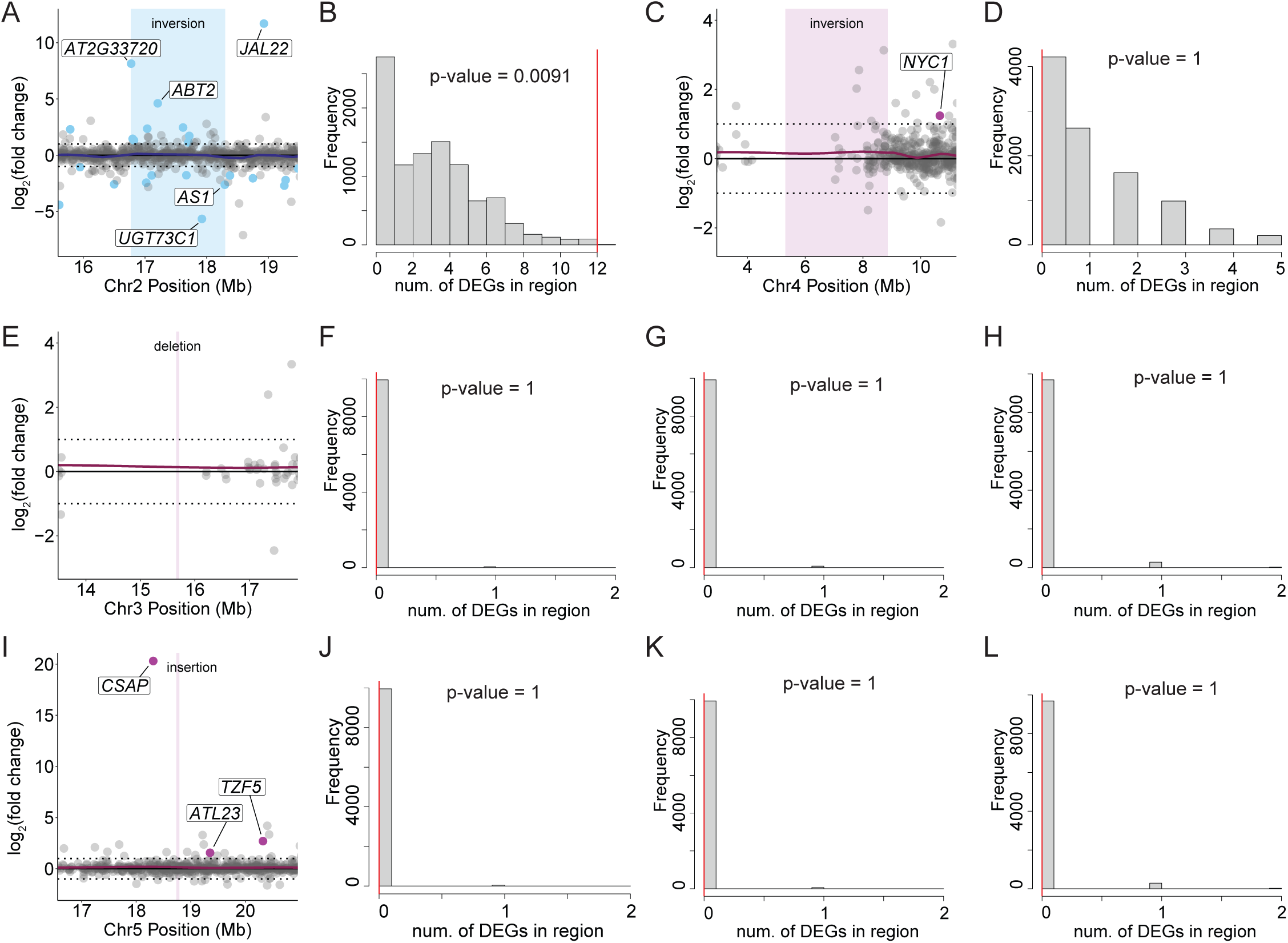
Permutation tests to determine if structural variants are associated with location of differentially expressed genes. **(A)** Log_2_(fold change) of genes within and 1 Mb outside the 1.52 Mb inversion in *BR-like dwarf.* DEGs are in blue and SVs are shaded in light blue. Dark blue line represents locally estimated scatterplot smoothing to aid pattern visualization. **(B)** Permutation test to determine if the 1.52 Mb inversion in *BR-like dwarf* plants contains more DEGs than expected by chance. All permutation tests were performed by selecting 10,000 genomic regions of an equivalent size to the region of interest at random. Histogram plots the distribution of number of DEGs contained within the randomly selected regions. Red line marks the number of DEGs in the region of interest. P-values calculated by the probability that permuted values exceed the observed value; if p-value is < 0.05 we conclude the region of interest is enriched for DEGs. **(C)** Log_2_(fold change) of genes within and 2 Mb outside the 3.54 Mb inversion in *short-internode dwarf.* DEGs are in pink and SVs are shaded in light pink. Dark pink line represents locally estimated scatterplot smoothing to aid pattern visualization. **(D)** Permutation test for 3.54 Mb inversion in *short-internode dwarf* plants. **(E)** Log_2_(fold change) of genes 2 Mb outside 176 bp deletion in *short-internode dwarf.* DEGs are in pink and SVs are shaded in light pink. Dark pink line represents locally estimated scatterplot smoothing to aid pattern visualization. Permutation tests for **(F)** 5 kb, **(G)** 10 kb, and **(H)** 50 kb flanking the deletion on Chr3 in *short-internode dwarf* plants. **(I)** Log_2_(fold change) of genes 2 Mb outside 184 bp insertion in *short-internode dwarf.* DEGs are in pink and SVs are shaded in light pink. Dark pink line represents locally estimated scatterplot smoothing to aid pattern visualization. Permutation tests for **(J)** 5 kb, **(K)** 10 kb, and **(L)** 50 kb flanking the insertion on Chr5 in *short-internode dwarf* plants.

Finally, we evaluated the relationships between structural variation identified by Nanopore sequencing and gene expression in *BR-like dwarf, short-internode dwarf,* and *variegated* mutants. Both sequenced *BR-like dwarf* plants contain a 1.52 Mb inversion on chromosome 2 that is not present in non-phenotypic control plants of the same line (Figure 4B,F, Table S11) and we investigated gene expression surrounding this locus further. The 1.52 Mb inversion was significantly enriched for differentially expressed genes, as determined by a permutation test (Figure S9A,B). Examining the inversion breakpoints more closely, the right breakend separated the first 42 bp of the 5’UTR of *ASYMMETRIC LEAVES 1 (AS1; AT2G37630*) from the rest of the gene (Figure 4F).

Near the left breakend of the inversion, the expression of an AP2/B3-like transcriptional factor family protein (AT2G33720) was increased in leaves (Figure 2A, Figure 4F, Table S3). Taken together, these data suggest that the 1.52 Mb inversion resulted in an altered *AS1*-mediated transcript abundance, influencing leaf patterning in *BR-like dwarf* plants.

Three SVs were common to the two *short-internode dwarf* plants that were sequenced via Nanopore: a 176 bp deletion on chromosome 3, a 3.54 Mb inversion on chromosome 4, and a 184 bp insertion on chromosome 5 (Figure 4B,C, Table S11). The large inversion on chromosome 4 was not enriched for DEGs, and neither were regions surrounding the deletion on chromosome 3 or the insertion on chromosome 5 (Figure S9C-L). However, since gene expression data was derived from only the leaves of *short-internode dwarf* plants, we cannot exclude the possibility that these structural variants have an impact on gene expression in other plant organs. No structural variants were identified in *virescent* plants utilizing either short- or long-read data. Although it is possible that no structural variation is present in this line, it is also possible that *virescent* plants contain an SV that cannot be detected with our methods due to its complexity or genomic location.

Long-read sequencing of the *variegated* line identified a 37 bp deletion in the third exon of *IM*; this caused a frame-shift resulting in a premature stop codon (Figure 4E). Crossing plants of the *variegated* phenotype to known *immutans* mutants failed to complement the variegated phenotype (32/32 variegated when crossed to *im-spotty* CS3639 [68], 34/34 variegated when crossed to *im-spotty* CS73029 [69] and 42/42 variegated when crossed to *im* CS3157), indicating that the 37 bp deletion in *IM* is indeed the causal locus for *variegation*.

In summary, our genomic analyses of mutants shows that etoposide treatment can create novel plant traits via the induction of SVs including inversions, deletions, duplications, and translocations.

## Discussion

We describe a simple and convenient technique that enables researchers or breeders with limited resources to induce structural variation. We found that early exposure of *A. thaliana* to the Topo II inhibitor etoposide induces heritable, large-effect mutations at high frequency. Examination of M2 plants found that 24/37 mutant lineages had at least one out of six assessed phenotypes. By short- and long-read genomic sequencing of mutagenized lines—including those that displayed novel phenotypes—we identified the full spectrum of structural variations. Analysis of short-read data identified 2.53 SVs per mutagenized M1 line whereas long-read data identified 3.25 SVs per mutagenized M1 line, with one event being detected in both short and long-read data. The lower number of SVs identified by short-read data is consistent with the poorer performance of short-read SV callers relative to callers using long-reads [64]. Our data suggest that mutation rate and type induced by etoposide exposure is comparable to mutant populations generated by irradiation. Analysis of Arabidopsis seeds irradiated with γ rays or carbon ions showed mutations occurring at a rate of 0.8-1.6 SVs per mutant lineage [70]. Thus, etoposide treatment out-performed irradiation by producing more than 2.5 events per mutagenized plant. A further increase in the number of mutations per M1 parent could perhaps be attained either by prolonging exposure to etoposide beyond two weeks or by germinating seeds on media containing a cocktail of etoposide and other drugs that induce genomic instability such as topoisomerase I inhibitors [71] or the DNA damage-inducing agent cisplatin [72].

Analysis of all detected SVs suggests that etoposide disproportionately impacts genes, although our analytical approaches cannot conclusively identify all SVs in heterochromatin. The SVs that we identified disrupt one or more genes simultaneously or create novel cis-regulatory regions. The effect of this genic bias is striking; even in our relatively small mutant population, multiple loss-of-function mutations were identified in a single line. This suggests that etoposide treatment can be used to create novel polygenic traits.

What explains this bias for mutations in genes? Topoisomerase II has been shown to associate with actively transcribed regions and open chromatin regions [42]. This association with actively transcribed regions suggests a future method to enrich for mutations in condition-specific genetic pathways by simultaneously exposing seeds to etoposide and another condition. For example, etoposide might promote mutagenesis of immunity genes whose transcription is triggered when seeds are germinated in the presence of microbial pathogens. In another scenario, seeds germinated in the presence of a hormone and etoposide might accumulate mutations in genes that respond to said hormone.

The inversions and translocations generated by this technique might also disrupt meiosis and distort segregation, but this possibility has not been tested. Etoposide also created duplications and deletions that could potentially alter gene dosage over mega-base scale regions. In allopolyploids, such segmental deletions can provide a deficiency chromosome that can be used to assess the relative contribution of homeologs to different traits.

How do we identify genetic changes that are causative for phenotypes of interest? Future studies will accelerate candidate-gene discovery by employing structural-variant callers, *de novo* genome assembly-based approaches, and RNA-seq based mapping [64]. Although our study did not aim to use RNA-seq to identify mutations, it provides an example of how RNA-seq data in tandem with genome sequencing can help shortlist potential causal mutations. In cases where a potential causative variant is not obvious, these strategies can be combined with traditional genetic mapping approaches. However, mapping-by-sequencing approaches might not be easily applicable to some classes of mutants—for example, inversions or translocations can suppress recombination and reduce the efficacy of mapping-by-sequencing.

In our current study, sequencing and close analysis of the genomes and transcriptomes of *BR-like dwarf, short-internode dwarf, virescent,* and *variegated* plants allowed us to link two of these phenotypes to strong candidate mutations. The *BR-like dwarf* phenotype is likely caused by an inversion that disrupts the expression of *ASYMMETRIC LEAVES 1* and the *variegated* phenotype is caused by a 37 bp deletion in *IMMUTANS.* Strikingly, nearly 60 years ago, the *asymmetric leaves* and *immutans* phenotypes were amongst the first X-ray induced mutations isolated in *A. thaliana* by Redei [73,74]. These results suggest that our easy-to-use chemical mutagenesis approach provides plant biologists and breeders with a new tool that can replace irradiation and reduce barriers to access tools necessary for the creation of structural variation.

## Materials and Methods

### Drug treatment

Etoposide (Abcam AB120227) was dissolved in DMSO to obtain a 100 mM solution that was added to 0.5x Murashige and Skoog (MS) media (supplemented with 2% sucrose and Phytoagar) to final concentrations of 0, 20, 40, 80, 160, 320, and 640 µM. We noticed decreased solubility of etoposide in aqueous solutions at concentrations beyond 160 µM. Wild-type Col-0 seeds were sterilized by incubating in 2% (v/v) PPM (Plant Cell Technologies) for three days at 4°C and plated on MS media containing etoposide or DMSO. Seeds were germinated and grown on this media for two weeks, and seedlings were examined for signs of stunted root or shoot growth. Two- to three-week-old seedlings (M1 plants), including several showing stunted growth, were transferred to soil and allowed to set seed. M2 seeds were collected from mutagenized M1 plants.

### Plant Growth Conditions

*Arabidopsis thaliana* was grown in a Conviron growth chamber at 22°C with 16 h of light per day (120 µmol).

### Phenotyping

Between 25 and 75 seeds from each M1 line were planted over three phases and M2 plants with visible phenotypes were manually identified. To identify mutants with leaf-shape differences, we measured leaf shape and size in mutant lineages where M2/M3 populations included three or more individuals displaying similar mutant phenotypes. Leaves were excised at the base of the petiole, arranged in decreasing size order on white paper, and scanned at 600 dpi on an Epson V600 scanner. Six automated leaf shape measurements were obtained (blade area, circularity, length, perimeter, width, and petiole length) using the LeafJ plugin for ImageJ software [75]. Leaf shape was manually traced in cases where leaf boundaries were not distinguished by ImageJ.

### RNA sequencing and analysis

RNA sequencing was conducted on M3 generation plants. Six plants of each *BR-like dwarf, short-internode dwarf*, and *virescent* phenotypes, as well as four non-phenotypic individuals corresponding to sibling lines of each phenotype were sequenced. For the fraction variegated line, thirteen green plants (three of which produced ∼25% variegated progeny) and one white plant were sampled. Nine M3 progeny of DMSO-treated plants were included as controls.

Mature leaf tissue from 32-day-old plants was flash frozen in liquid nitrogen. Frozen tissue was ground into a powder with a Tissuelyser II (QIAGEN; 30 beats/sec for 30 sec). RNA was extracted using an RNeasy Plant Mini Kit (QIAGEN). Library preparation was performed at the MIT BioMicro Center using an NEBNext Ultra II Directional RNA Library Prep Kit for Illumina (New England Biolabs). Samples were multiplexed on one lane of an Illumina NovaSeq 6000 S4 flow cell and sequenced at the MIT BioMicro Center with 150 bp paired end reads.

Trim Galore [76] was used to remove adapters and low-quality ends from reads (*--paired --retain_unpaired -q 25*). Salmon [77] was used to quantify the expression of transcripts, utilizing the Araport11 transcriptome as the reference and the Col-CEN v1.2 genome [65] as decoy sequences. Five samples were identified as outliers and removed from the dataset, based upon the PCAgrid function in the R package rrcov [78,79] and preliminary analyses. DESeq2 [80] was used to quantify differential expression; for a gene to be considered it must have at least 10 normalized counts in four or more samples. Each phenotype of interest was analyzed independently; DESeq2 design (*∼ treatment + phenotype*) incorporated both treatment (etoposide or DMSO) and phenotype (present or absent). The R package clusterProfiler [81] was used to conduct gene set enrichment analysis. Data was visualized utilizing R and packages ggplot2, EnhancedVolcano [82], and pheatmap [83].

### Illumina sequencing and identification of structural variants

DNA was extracted from the rosette leaves of M2 and M3 plants using a CTAB protocol. Up to 15 ng of genomic DNA was used by the BioMicro Center at MIT to construct DNA sequencing libraries using the Nextera DNA Flex library preparation kit. Libraries were multiplexed and sequenced in 150 bp paired-end format using an Illumina NovaSeq 6000 S4. The first 10 bp of each read as well as any adaptor sequence was removed using Trimmomatic [84] and aligned to the Col-0 Centromere assembly [65] using BWA-MEM [85] (*bwa mem -R "@RG\tID:id\tSM:sample\tLB:lib" reference.fasta sample.R1.fastq sample.R2.fastq \ | samblaster --excludeDups --addMateTags -- maxSplitCount 2 --minNonOverlap 20*). These alignments were used to identify structural variants by two separate methods. Larger deletions and duplications were identified using a method proposed by the Comai Lab [61,62]. Coverage was calculated for each 100 kb window for properly paired and aligned reads using Bedtools [86] (*bedtools coverage -a 100kb_windows.bed -b sample_paired_sorted.bam -counts*) and read-counts/window for each line was normalized to library size as reads per million.

The median normalized read-counts for all lines for each window was calculated. As most lines are not expected to share a structural variant, this median read depth is considered to be the wild-type genome dosage and is set to a ploidy or chromosomal dosage of two. The chromosomal dosage of any window for a given line is then calculated as (*normalized read-counts / median read-counts) x2.* Windows with a coverage score ≥3 were considered heterozygous duplications, ≥4 to be homozygous duplications, ≤1 to be heterozygous deletion and ≤0 to be a homozygous deletion.

In addition, we also identified structural variation using a benchmarked Lumpy Express pipeline [12,63]. Briefly, Samtools [87,88] was used to identify discordant and split reads from the SAM file that was output by BWA [85]. Files containing discordant reads, split reads, and all reads were used by Lumpy Express to identify putative structural variants. Due to mapping artifacts, Lumpy Express and other variant callers often identify false-positive structural variant candidates at repetitive regions [89]. To address this problem, we filtered out variants from ChrC, ChrM, and the NOR on chromosome 2. To remove SVs present in lab wild-type Col-0 stocks, we removed SVs within 2 kb of any SV found in lines only treated with DMSO, or SVs shared by four or more unrelated etoposide-treated mutant lineages. Finally, only those deletions and duplications that were supported by split-reads were retained in the final list of SVs.

To assess if etoposide causes single nucleotide variants, we used BWA-MEM to align reads from 32 M2/M3 progeny of etoposide-treated plants and four M2 progeny of control plants treated with DMSO to the Col-0 centromere assembly [65]. Aligned reads were sorted and converted to variant calling format (*bcftools mpileup -O b -o file.bcf -f ref.fasta file.sorted.bam*) followed by SNP calling (*bcftools call --ploidy 2 -m -v -o file.vcf file.bcf*) [87]. Variants that were called by bcftools were filtered to retain only calls with a quality score higher than or equal to 40 (*bcftools view -O z -o filtered.vcf.gz -e ‘QUAL<=40’ in.vcf.gz*). Variants shared by all untreated lines were extracted (*bcftools isec -n=5 -p common_positions.vcf input_files_filtered.vcf.gz*), and then excluded from all other treated lines (*bcftools view -T common_positions.vcf filtered_files.vcf -O z -o untreated_filtered_files.vcf*).

### Nanopore sequencing and identification of structural variants

Two mature leaves from 32-day-old M3 plants were flash frozen in liquid nitrogen. Frozen tissue was ground into a powder with a Tissuelyser II (QIAGEN; 30 beats/sec for 1 min). High molecular weight DNA was extracted using a Wizard HMW DNA Extraction Kit (Promega). DNA was sheared by passing through a 23G needle twice.

Contaminating RNA and proteins were removed using RNase A (New England Biolabs) and Proteinase K (New England Biolabs). DNA was further cleaned and small fragments removed using a modified SPRI bead mixture in which the buffer of Agencourt AMPure XP beads was exchanged with a custom buffer of 10 mM Tris-HCl, 1 mM EDTA pH 8, 1.6 M NaCl, 11% w/v PEG 8000 [90]. To perform size selection, in brief, 0.7x volume of modified bead solution was added to each DNA sample, then incubated on a nutator for 10 min. Samples were placed on a magnetic stand to bind the beads to the side of the tube, then washed twice with 70% ethanol. Beads were not allowed to dry before adding nuclease-free water to elute the DNA. To increase yield and remove residual ethanol, samples were incubated at 50°C for 2 min and then at room temperature for two hours with the lids open.

All library preparation and sequencing was performed at the MIT BioMicro Center. For two samples showing the *BR-like dwarf* phenotype, high molecular weight DNA extractions had low yield. For these samples and corresponding controls (individual from sibling line without dwarf phenotype and individual from a line treated with DMSO), library preparation was performed utilizing a Rapid Barcoding Kit V10 (Oxford Nanopore Technologies; SQK-RBK110-96). These four samples were multiplexed and sequenced on a single PromethION R9.4.1 flow cell (Oxford Nanopore Technologies; FLO-PRO002). For all other samples, library preparation was performed with a Native Barcoding Kit V14 (Oxford Nanopore Technologies; SQK-NBD114-24). These samples were multiplexed into groups of three and each group was sequenced on its own PromethION R10.4.1 flow cell (Oxford Nanopore Technologies; FLO-PRO114M). Both runs utilized the following software versions for sequencing and base-calling: MinKNOW 23.04.6, Bream 7.5.10, Configuration 5.5.14, Guppy 6.5.7, and MinKNOW Core 5.5.5.

Nanopore reads were filtered utilizing Filtlong v0.2.1 [91] to select the best reads in terms of length and quality, up to a max coverage of 50x (*--min_length 1000 -- min_window_q 40 --trim --split 1500 --target_bases 6605000000*). Filtered reads were mapped to the Col-CEN v1.2 genome [65] using Vulcan dual-mode alignment pipeline [92], which leverages both minimap2 [93] and NGMLR [94], using default settings for ONT reads. Structural variants in each sample were detected with cuteSV *(--max_cluster_bias_INS 100 --diff_ratio_merging_INS 0.3 --max_cluster_bias_DEL 100 -- diff_ratio_merging_DEL 0.3 --min_size 30 --max_size -1 –genotype;* [66,67]). SVs identified in all samples were merged with SURVIVOR (*merge 1000 1 1 1 0 30;* [95]), then samples were genotyped with cuteSV for all SVs identified (*--min_mapq 20 -- max_cluster_bias_INS 100 --diff_ratio_merging_INS 0.3 --max_cluster_bias_DEL 100 -- diff_ratio_merging_DEL 0.3 --min_size 30 --max_size -1*). Structural variants present in the DMSO and/or Col-0 controls, or present in more than one phenotypic line, or with a quality less than 20 were removed. Data was visualized in R using ggplot2.

## Data Availability

Short-read DNA sequencing, long-read DNA sequencing, and mRNA-seq data are available in GEO records GSE284455, GSE284456 and GSE284457.

## Declaration of Interest

Whitehead Institute has filed a patent application related to this work.

## Acknowledgements

We thank Aysha Hussain, Jaza Alam, and Lingfeng Shi for assistance with phenotyping, Dyna Louis for assistance with verifying structural variants, and Xinlei Gao for assistance with statistical testing. This research was funded by grants to MG from The Abdul Latif Jameel Water and Food Systems Lab at the Massachusetts Institute of Technology, from the Professor Amar G. Bose Research Grants, and from a gift to MG from the Dr. Vincent J. Ryan Orphan Plant Project. MG is an Investigator of the Howard Hughes Medical Institute.

## Supplemental Information

Table S1: List of phenotypes observed in M2 progeny of etoposide-treated plants. Table S2: List of RNA-Seq libraries.

Table S3: DESeq2 output identifying mis-regulated genes in *BR-like dwarf*.

Table S4: DESeq2 output identifying mis-regulated genes in *short-internode dwarf*.

Table S5: DESeq2 output identifying mis-regulated genes in virescent. Table S6: List of samples sequenced with Illumina.

Table S7: Relationship between the sequenced lines, treatments, and nature of sequencing.

Table S8: Deletions and duplications identified by read coverage analysis.

Table S9: Structural variants detected by LUMPY Express using short-read sequencing.

Table S10: List of samples sequenced with Nanopore.

Table S11: Structural variants identified using Nanopore long-read data.

## Notes

https://www.ncbi.nlm.nih.gov/geo/query/acc.cgi?acc=GSE284455

https://www.ncbi.nlm.nih.gov/geo/query/acc.cgi?acc=GSE284456

https://www.ncbi.nlm.nih.gov/geo/query/acc.cgi?acc=GSE284457

### Summary of Updates

Reviewer suggestions have been incorporated into the manuscript. Figure 3 and 4, and associated text have been updated.

